# Single-cell metabolic imaging and digital scoring of fat tissue remodeling by label-free metabolic microscopy

**DOI:** 10.1101/2025.06.06.658032

**Authors:** Myeongseop Kim, Constantin Berger, Alexander Wolf, Alina Peteranderl, Martin Klingenspor, Vasilis Ntziachristos, Yongguo Li, Miguel A. Pleitez

## Abstract

Adipose tissue plasticity and functional heterogeneity play a central role in maintaining energy homeostasis, and their malfunction leads to metabolic disorders such as obesity, diabetes, and cardiometabolic disease. Rapid, single-cell metabolic imaging of intact fat tissue not only extends our understanding of metabolic dynamics and heterogeneity but also holds great potential as a tool for clinical diagnosis. However, the use of exogenous labels and dyes in conventional optical microscopy results in tissue deformation and requires time-consuming tissue preparation. Here, we demonstrated single-cell imaging of metabolic changes and heterogeneity in freshly excised adipose tissues that can distinguish tissue types without the need for exogenous labels using bond-specific, non-destructive, mid-infrared optoacoustic microscopy (MiROM) that allows preserving the native tissue architecture with minimal sample preparation time. Further leveraging MiROM, we monitored intracellular molecular and morphological changes during postnatal remodeling of adipose tissue when metabolic characteristics of adipocytes undergo a transient drastic change. Additionally, we developed an AI-based quantitative spatial tissue analysis tool (Q-SAT) to predict the spatial distribution of white fat- and brown fat-like features, providing a robust digital scoring method for adipose tissue phenotypic assessment. Collectively, we implemented MiROM as an enabling technology to provide fast, label-free metabolic imaging of unprocessed adipose tissue, opening a new perspective for understanding and characterizing the morpho-functional dynamics of adipose tissue remodeling.

## Introduction

Adipose tissue is an extraordinarily plastic and heterogeneous organ in mammals that controls energy balance, glucose, and lipid metabolism^1–8^ and is commonly classified into two types: white adipose tissue (WAT), which stores energy, and brown adipose tissue (BAT), which dissipates energy and contributes to thermogenesis and metabolic control. The remodeling capacity of adipose tissue not only allows the body to adapt rapidly to variations in energy supply and demand but also plays a major role in orchestrating metabolic health.^1^ The remarkable plasticity of adipose tissue is further exemplified by the ability of adipocytes to interconvert between white and brown phenotypes. Brown-like adipocytes or brite (brown-in-white, or also beige, or recruitable brown) adipocytes share the capacity for uncoupling protein 1 (UCP1)-mediated non-shivering thermogenesis yet present a discrete cell type. These beige adipocytes can be found interspersed within WAT, demonstrating the complexity of fat depots and the need for understanding not only its plasticity but also heterogeneity.^9,10^ As impaired plasticity and loss of heterogeneity lead to diminished or abnormal responses to physiological challenges and thus drive the progression of metabolic diseases, a comprehensive understanding of adipose tissue plasticity and heterogeneity is crucial for developing new potential therapeutic targets for managing obesity and other metabolic disorders, ultimately advancing relevant clinical applications.

Metabolic imaging allows the acquisition of spatio-temporal information on the biological response to external stimuli. In cell and animal models, it can provide key information on the progression of a disease as well as on the performance of a particular drug or treatment. However, most of the technologies available for metabolic imaging are restricted to being used in combination with exogenous labels or markers that have limited biodistribution, can potentially perturb metabolic responses, or are simply not available. Moreover, metabolic responses can be highly dynamic and heterogeneous—both temporally and spatially. This highlights the necessity to study the metabolic plasticity with high sensitivity and even at single-cell resolution. Conversely, conventional technologies used for adipose tissue analysis often perform only bulk measurements that ignore the heterogeneity of cell populations and cell-to-cell variations. Additionally, although state-of-the-art single-cell RNA-Seq and spatial transcriptomics can reveal cellular heterogeneity at the transcriptomic level, they are not able to provide a direct quantitative assessment of metabolic response because they rely on gene expression patterns to indirectly infer metabolic activity. Therefore, new technologies that enable a holistic understanding of the metabolic plasticity and functional heterogeneity at a single-cell level that provide complementary insights into the underlying biological mechanisms are highly demanded.

Label-free chemical imaging techniques, unlike imaging with exogenous labels, enable the analysis of morphological characteristics and molecular information of tissue while preserving the native architecture of the tissue of interest and avoiding the time-consuming sample preparation steps typically required for exogenous staining. In particular, vibrational spectroscopic methods based on Raman scattering^11,12^ or mid-infrared (mid-IR) absorption^13,14^ have been developed for the label-free chemical imaging. Raman-scattering microscopy modalities, i.e., spontaneous Raman scattering, coherent anti-Stokes Raman scattering (CARS) and stimulated Raman scattering (SRS) microscopy, have been applied in cell and tissue studies to characterize and to monitor the various biomolecules (e.g., lipid and protein) during metabolic activities.^12,15^ For instance, Raman-scattering microscopy has recently revealed distinct cellular phenotypes of white and brown adipocytes during adipogenic differentiation.^16,17^ However, the low Raman scattering cross-sections of biomolecules—typically on the level of 10^−30^ cm^2^—result in low detection sensitivity (low mM range) and high irradiation energies (100’s of mW) on the studied specimens, which can lead to sample perturbation by photodamage.^18^ On the contrary, mid-IR absorption cross-sections are up to eight orders of magnitude larger than Raman-scattering^19–21^ and thus mid-IR spectroscopy and imaging offer the potential to significantly improve sensitivity and specificity for label-free analytical histology in the mid-IR region (4000-400 cm^−1^).^22^ However, as conventional mid-IR spectroscopy relies primarily on optical detection, the applicability of mid-IR microscopy on living cells and unprocessed tissues is limited by the significant IR absorption of water and signal loss caused by negative-contrast optical detection, i.e., the detected signal becomes weak as optical absorption increases.^23,24^ As a result, assessment of excised tissues by optically-detected mid-IR microscopy requires time-consuming tissue preparation steps similar to those required in conventional histology, i.e., tissue sectioning slices <10 μm thickness.^23^

Unlike conventional vibrational microscopy, mid-IR optoacoustic microscopy (MiROM) uses detection of optically generated acoustic signals—i.e., optoacoustic (OA) signals—which are less attenuated and scattered than purely optical signals. MiROM enables obtaining mid-IR absorption contrast from thick, unprocessed, freshly-excised tissues (up to 6 mm thickness) with minimal preparation steps^25^—thus preserving the integrity of unprocessed tissue during histological analysis while providing high chemical sensitivity. MiROM has been applied to the histological characterization of carotid atherosclerosis,^26^ which visualized high-risk plaque features with molecular assignment. MiROM has also been applied for label-free differentiation between inflamed vs. non-inflamed WAT^27^ based on spectroscopic feature selection, achieving a considerable reduction of total imaging time (by a 15x factor) as compared to laser scanning confocal microscopy, which uses labels for identifying structures associated with inflammation.^27^ Furthermore, MiROM has demonstrated depth-selective analysis of OA signal applied to minimize the interference of superficial skin layers to enhance the sensitivity for non-invasive glucose sensing.^28^

Here, we took advantage of the high heterogeneity and plasticity of adipose tissue as a model system and hypothesized that the depth-selectivity and label-free chemical specificity imaging features of MiROM make it possible to identify the intrinsic hallmarks of WAT and BAT in fresh unprocessed fat tissues. We found that, with minimal sample preparation, MiROM allowed us to identify intrinsic biochemical characteristics that can distinguish BAT from WAT and monitor the transient intracellular behavior of white fat depots observed during the postnatal development of mice. Specifically, in this work, for the first time, we characterized the spatial-temporal distribution of lipid and protein changes in beige adipocytes at the cellular level during postnatal tissue remodeling, i.e., browning and whitening. Additionally, we leveraged the ability to distinguish spectral and morphological characteristics between WAT and BAT by developing a tailored AI-aided quantitative spatial tissue analysis tool (Q-SAT), which allows us to digitally score the spatial distribution of tissue features corresponding to WAT and BAT (WAT&BAT scoring). Overall, MiROM combined with Q-SAT provides a robust biotechnology for label-free histological assessments of adipose tissue, allowing characterization and aiding in understanding the morpho-functional dynamics of adipose tissue remodeling.

## Results

### Intracellular imaging and multi-spectral analysis of adipocytes in freshly excised fat tissue

MiROM is a label-free chemical imaging technique that preserves the native architecture and intrinsic biomolecular composition of freshly excised tissues and thus enables circumventing the limitations of conventional methods for histological assays (e.g., H&E staining, immunohistochemical staining). To demonstrate MiROM’s capabilities for histological assays, we implemented it for label-free chemical analysis of freshly excised adipose tissue as illustrated in **Fig.1a**; details on MiROM operation are given in **Methods** and elsewhere.^25–28^ In short, in MiROM, an ultrasound (US) signal is generated by the optical absorption of a diffraction-limited mid-IR excitation beam (∼5 μm focus size at 2850 cm^−1^ excitation) illuminating the tissue and detected by a focused ultrasound transducer. The focused mid-IR excitation beam—from a broadly tunable quantum cascade laser (QCL)—and a focused US detector are confocally aligned to the adipose tissue that is placed on a custom-designed Petri dish with deionized water for acoustic coupling between the tissue and the US detector. In this configuration, mid-IR spectra on selected structures of interest in the tissue can be obtained by tuning the excitation wavelength of the QCL across a broad spectral range (2932-909 cm^−1^). **Fig.1b** shows a representative mid-IR OA spectrum obtained from white adipocytes. As expected, since lipid droplets in adipocytes are primarily composed of triglycerides,^29^ the spectrum in **Fig.1b** is dominated by the spectral features of triglycerides, including symmetric CH_2_ stretching (2856 cm^−1^), C = C stretching (1650 cm^−1^), and C - O stretching vibration of the C-OH group (1116 cm^−1^).^22,30^ Next, a MiROM micrograph is obtained by point-by-point raster scanning the sample across the focal plane while acquiring the OA signal at a selected mid-IR excitation wavelength (**Supplementary Fig.1a, b**) and plotting the maximum amplitude projection (MAP) of the OA signal for each measured point. Visualization of the different intrinsic bio-molecular content of tissues is obtained by acquiring micrographs at multiple vibrational transitions (mid-IR excitation wavelengths). For instance, as shown in **Supplementary Fig.1a, b**, lipid and protein content of epididymal white adipose tissue (eWAT) and interscapular brown adipose tissue (iBAT) is visualized at 2856 and 1550 cm^−1^, respectively. Here, we observe that, although an overlap from different molecular content is expected, the contrast obtained for micrographs at 2856 cm^−1^is greatly dominated by the lipid content in adipocytes, i.e., the triglycerides in lipid droplets, while the contrast obtained at 1550 cm^−1^ mainly originates from the protein and water content in the extracellular matrix (ECM) with only a small contribution from the interior of adipocytes. As a result, since the ECM is composed of lower lipid and higher protein content compared to adipocytes and vice versa, we observed a considerable contrast difference between adipocytes and extracellular protein structure obtained at 2856 and 1550 cm^−1^, and thus micrographs at these wavelengths appear to be complementary, see **Supplementary Fig.1a, b**.^31–33^ At the location of adipocytes, due to adipocytes’ low protein and water content, micrographs at 1550 cm^−1^appear dark compared to the ECM. However, although rather low, intracellular protein and water contrast at 1550 cm^−1^ is still expected from adipocytes, but we hypothesized that it is overshadowed by the strong contrast of the surrounding ECM and thus not visible at first sight in the micrographs. This hypothesis was tested and proven positive by closer inspection of the OA signal from these seemingly dark areas, where we found non-negligible contrast content at the adipocytes’ location, see **Supplementary Fig.1g, h**.

**Figure 1.**
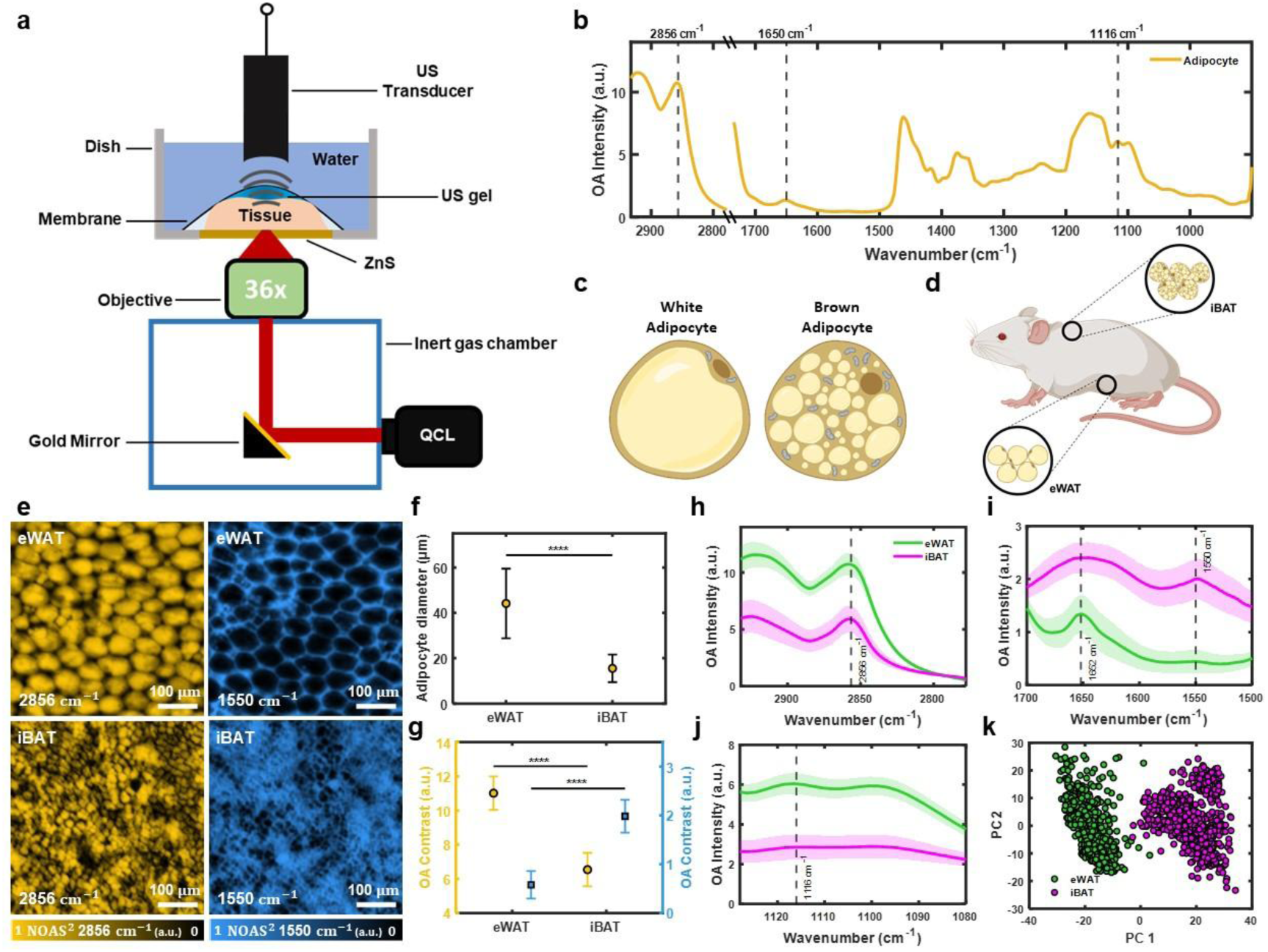
Determination of spectral and morphological hallmarks of white and brown adipose tissue. **(a)** Schematic representation of mid-infrared optoacoustic microscopy (**MiROM**) for label-free analytical histology of adipose tissue. Ultrasound (**US**), quantum cascade laser (**QCL**). **(b)** Representative optoacoustic (**OA**) spectra (2932-909 *cm*^−1^) from a selected adipocytes in adipose tissue. **(c)** Illustration of a white adipocyte (left) and a brown adipocyte (right). **(d)** Schematic diagram of adipocyte tissue locations on FVB/NJ mice. The locations of epididymal white adipose tissue (**eWAT**) and interscapular brown adipose tissue (**iBAT**) are marked. **(e)** Depth-selective OA micrographs—Field-Of-View (**FOV**) of 1 × 1 *mm*^2^ —of eWAT and iBAT for 2856 *cm*^−1^ (yellow) and 1550 *cm*^−1^ (cyan). Normalized optoacoustic signal (**NOAS**). **(f)** Comparison of mean diameter of adipocytes in eWAT and iBAT. The error bars indicate standard deviation. **(g)** Comparison of digitally isolated adipocyte OA contrast for 2856 and 1550 *cm*^−1^ micrographs obtained from eWAT and iBAT adipocytes. The error bars indicate standard deviation. Asterisks (****) denote the level of statistical significance (P<0.0001) analyzed by a two-sample t-test. **(h-j)** Depth-selective OA spectra of eWAT and iBAT adipocytes for three different spectral ranges (**h**: 2932-2778 *cm*^−1^, **i**: 1700-1500 *cm*^−1^, and **j**: 1128-1080 *cm*^−1^). The mean spectra (solid line) and their standard deviations (shaded area) are shown for eWAT and iBAT. **(k)** Principal component (**PC**) analysis 2D scatter plot derived from eWAT and iBAT spectra. Results in **f-k** show seven independent measurements (N=7) for each tissue type. Results in **h-k** were obtained from over 100 adipocytes per mice. Illustrations **c, d** were created with BioRender.com.

Having demonstrated non-negligible contrast in adipocytes at 1550 cm^−1^ and in order to minimize the overshadowing effect of the surrounding ECM to precisely assess the intracellular content of adipocytes, we performed a digital isolation protocol of adipocyte contrast in two steps. First, we leveraged the depth-selective abilities of MiROM to reach the inner content of adipocytes by analyzing OA signals at selected depths. Depth-selective mid-IR spectral analysis is possible because the time-resolved (transient) OA signal contains depth information as reported by the arrival time of the ultrasound signal at the transducer—namely, signals from deeper tissue layers arrive with a time delay compared to signals from shallow depths. In this way, depth-selective information can be retrieved (axial isolation) by time gating the OA transient signals, see **Methods**, and **Supplementary Fig.1c-f** for details. Second, we applied an intensity threshold to the micrographs at 2850 cm^−1^ to generate a mask for 2D localization of the adipocyte-containing areas (lateral isolation).^28^

In the following, we analyzed the intrinsic intracellular content of adipocytes in adipose tissues by axial and lateral isolation.

### Determination of the morphological and molecular hallmarks of WAT and BAT by MiROM

The functional differences between WAT and BAT have been closely associated with the morphological differences of the adipocytes composing these two tissue types; for instance, adipocyte size, adipocyte density (count per unit area in mm^2^), and number of mitochondria as depicted in **Fig.1c**.^34,35^ In mice, epididymal white adipose tissue (eWAT) is representative for classical WAT, while interscapular brown adipose tissue (iBAT) is representative for classical BAT (**Fig.1d**).^7,36^ Based on strong contrast between adipocytes and extracellular protein structure observed at 2856 cm^−1^, we utilized MiROM to characterize morphological differences in adipocytes between eWAT and iBAT. Specifically, we quantified adipocyte size (diameter) and adipocyte density in both tissue types using samples from 14 mice (eWAT: N=7, iBAT: N=7). Representative depth-selective OA micrographs acquired at 2856 and 1550 cm^−1^ for eWAT and iBAT are shown in **Fig.1e** revealing a significant difference in adipocyte size between the two tissue types (**Fig.1f**, P≈0). Specifically, the diameter of adipocytes in eWAT was, on average, ca. 2.8 times larger than those in iBAT (eWAT: 44±15 μm, iBAT: 16±6 μm) while the adipocyte density in eWAT was ca. 8.5 times lower than in iBAT (**Supplementary Fig.2e, f**).

Moreover, by digital isolation of adipocyte contrast—i.e., depth-selected and laterally segmented contrast—we were able to compare eWAT and iBAT adipocyte-specific molecular content at 2856 and 1550 cm^−1^ (see **Fig.1g**). Here we found that, at 2856 cm^−1^ the contrast in eWAT is ca. 1.7 times higher than in iBAT while at 1550 cm^−1^ the contrast of iBAT is ca. 3.4 times higher than eWAT. Thus, the vibrational contrast information indicates a lipid-rich and protein-poor content for eWAT and a lipid-poor and protein-rich content in iBAT. These remarkable morphological and contrast differences between eWAT and iBAT observed at 2856 cm^−1^ (lipid) and 1550 cm^−1^ (protein) represent unique tissue-specific hallmarks that we used for tissue identification.

Beyond morphology, MiROM was also able to reveal remarkable spectral differences between eWAT- and iBAT-adipocytes in the 2932-909 cm^−1^ spectral range, see **Fig.1h-j** (for a total of 14 independent measurements, N=14). In particular, spectral differences between eWAT and iBAT were detected at the CH_2_ vibration region (2932-2778 cm^−1^; **Fig.1h**), the amide I and II bands (amide I band: 1700-1600 cm^−1^, amide II band: 1600-1500 cm^−1^; **Fig.1i**), and the CO vibration region (1128-1080 cm^−1^; **Fig.1j**). While the CH_2_ vibrational region (2932-2778 cm^−1^; **Fig.1h**) and the CO vibrational region (1128-1080 cm^−1^; **Fig.1j**) are relevant for lipid characterization, the amide I and II bands (1700-1500 cm^−1^) are critical to identifying protein content. From the CH_2_ and the CO vibrational regions—as reported by the area under the curve (AUC), see **Methods**—we observed ca. 2 times higher OA signal for eWAT-adipocytes compared to the OA spectra of iBAT-adipocytes. Spectral intensity differences in these two regions support the hypothesis that eWAT-adipocytes have a higher lipid content than iBAT-adipocytes. From the amide I and II bands, we obtained ca. 2.8 times higher AUC of mean spectra for iBAT-adipocytes than for eWAT-adipocytes suggesting that iBAT-adipocytes contain more protein than eWAT-adipocytes—most likely, due to the elevated mitochondrial count in iBAT-adipocytes as compared to eWAT-adipocytes. In particular, from the spectral region between 1700 and 1500 cm^−1^, we observed the most remarkable spectral differences between eWAT- and iBAT-adipocytes. In iBAT, we observed the two typical absorption bands characteristic of protein content at 1650 and 1550 cm^−1^ (amide I and II, respectively), while for eWAT, we observed only a narrow absorption band at 1650 cm^−1^ (no band at 1550 cm^−1^) with a slope towards 1700 cm^−1^ mostly associated with lipid content. The spectral differences between eWAT- and iBAT-adipocytes (for a total of 14 independent measurements, N=14) are visualized in **Fig.1k** where a 2D scatter plot obtained by principal component analysis (PCA) shows two clearly defined clusters grouped by spectra similarity, each group representing eWAT- or iBAT-adipocytes. Thus, MiROM is able to morphologically and spectrally distinguish between brown and white adipose tissues.

### Label-free longitudinal monitoring of postnatal adipose tissue remodeling

Besides classical white and brown adipocytes, brown-like adipocytes (brown-in-white, or also beige, or recruitable brown) occurring within WAT under certain conditions (e.g., cold exposure, beta-adrenergic stimulation, PPARγ agonist treatment) represent a distinct type of thermogenic fat cell and hold great promise for promoting metabolic health.^1,7,10^ Remarkably, during the first two months of postnatal development, iWAT of mice undergoes a drastic, but transient, remodeling process between a white-to-brown (browning – postnatal weeks 2-4) and brown-to-white phenotype (whitening – postnatal weeks 4-6) with unclear molecular control and physiological function (**Supplementary Fig.4**).^3,7,36,37^ We leveraged this dynamic postnatal adipose tissue remodeling in iWAT as an excellent model to test the robustness of MiROM in monitoring intracellular molecular and morphological changes of browning and whitening, longitudinally. Based on the morphological and molecular hallmarks of WAT and BAT identified using MiROM, we performed longitudinal monitoring of adipose tissue remodeling during postnatal development of mice (**Fig.2b**). **Fig.2a** shows the locations where tissues were excised as well as the region of interest (ROI) beneath the lymph node in iWAT where significant beige adipocyte development occurs. As expected, from OA micrographs (**Fig.2c**, **d**), we observed a decrease in the size of adipocytes in iWAT during postnatal weeks 2-4 and an increase in the size of adipocytes in iWAT during postnatal weeks 4-6 (**Supplementary Fig.2d, e**). In contrast, adipocyte count per unit area exhibited the opposite trend (**Supplementary Fig.2f**). The observed changes indicate adipocyte hyperplasia from weeks 2 to 4 and hypertrophy from weeks 4 to 6. These morphological changes were confirmed by hematoxylin and eosin (H&E) staining of the tissues (**Supplementary Fig.4**).

**Figure 2.**
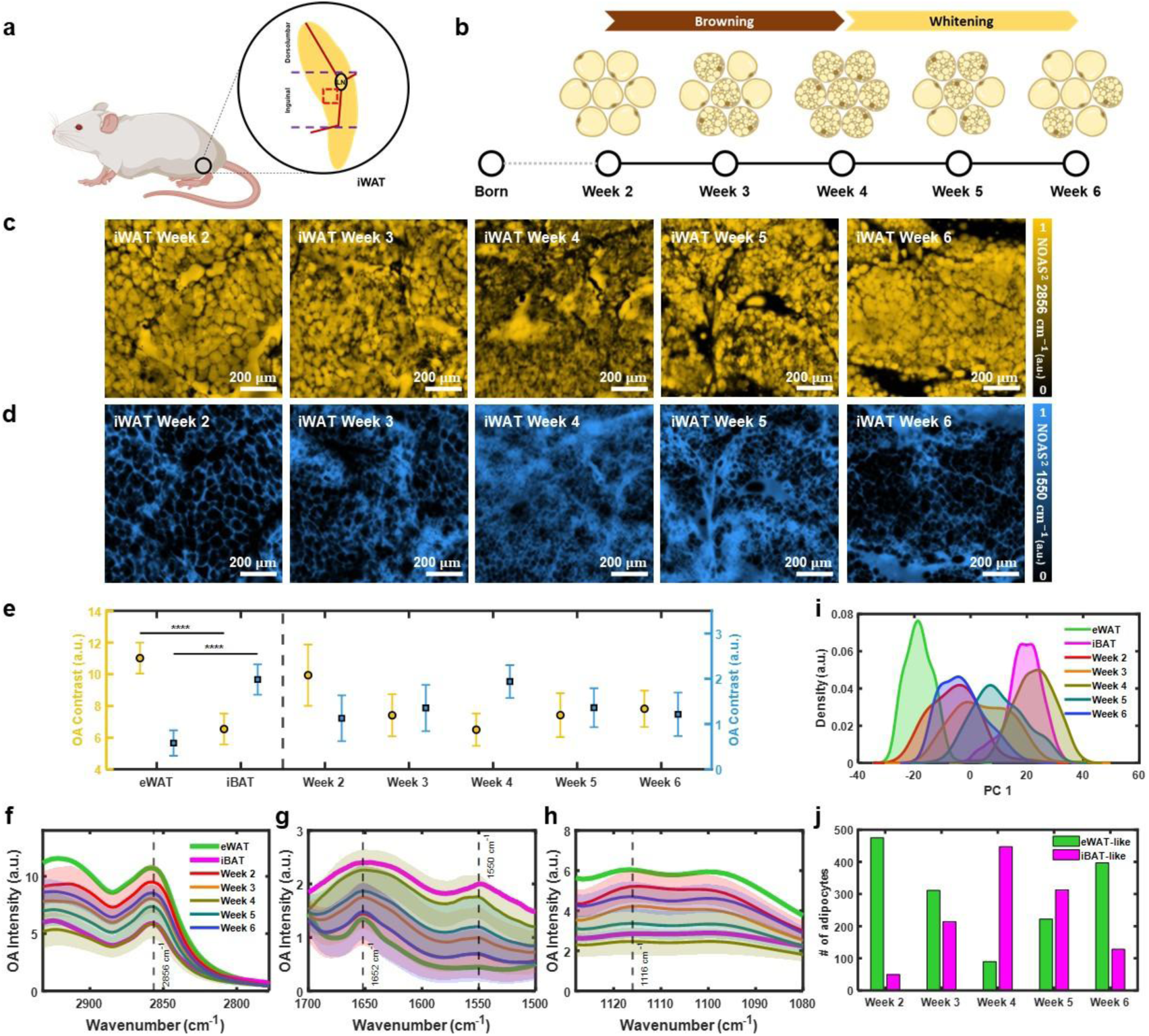
Monitoring morphological and spectral dynamics during postnatal development in inguinal white adipose tissue. **(a)** Schematic diagram of inguinal white adipose tissue (**iWAT**) location on FVB/NJ mice. The region of interest for iWAT, located beneath the lymph node (**LN**), is marked by a red dashed box. **(b)** Timeline of iWAT transition from postnatal week 2 through week 6 indicating a browning phase during postnatal weeks 2-4 and a subsequent whitening phase from postnatal weeks 4-6. **(c, d)** Representative example of depth-selective OA micrographs of iWAT at postnatal weeks 2-6 at 2856 *cm*^−1^ (c) and 1550 *cm*^−1^ (d) for a FOV of 1×1 *mm*^2^. **(e)** Comparison of digitally isolated adipocyte OA contrast (at 2856 and 1550 *cm*^−1^) for eWAT and iBAT (N=7, for each tissue type) with iWAT adipocyte contrast during postnatal weeks 2-6 (N=5, for each tissue type). The error bars indicate standard deviation. **(f-h)** Change of depth-selective OA spectra of iWAT adipocytes from postnatal weeks 2-6 for three different spectral ranges (**f**: 2932-2778 *cm*^−1^, **g**: 1700-1500 *cm*^−1^, and **h**: 1128-1080 *cm*^−1^). The mean spectra (solid line) and their standard deviations (shaded area) are shown for eWAT and iBAT (ca. 100 adipocytes per mice, N=7), and iWAT (ca. 100 adipocytes per mice per postnatal week, N=5). **(i)** Kernel density estimation of PC1 derived from adipocyte spectra for eWAT, iBAT, and iWAT (postnatal weeks 2-6). **(j)** Bar chart indicating the estimated amount of eWAT-like and iBAT-like adipocytes in iWAT during postnatal weeks 2-6 obtained by a logistic regression model trained on spectra from eWAT and iBAT adipocytes. Illustrations **a, b** were created with BioRender.com.

For a thorough comparison, vibrational contrast of eWAT, iBAT, and iWAT (eWAT and iBAT: N=7; iWAT at postnatal weeks 2-6: N=5 each) at two channels (lipid: 2856 cm^−1^; protein: 1550 cm^−1^) is shown in **Fig.2e**. The contrast at 2856 cm^−1^ decreases in iWAT from postnatal weeks 2 to 4 and increases again from postnatal weeks 4 to 6, while the contrast at 1550 cm^−1^follows opposite trends. These observations match the transient browning (postnatal weeks 2-4) and subsequent re-whitening (postnatal weeks 4-6) phases of iWAT, as confirmed by western blot and whole-mount immunofluorescence staining (**Supplementary Fig.4**).

**Fig.2f-h** shows depth-selective OA spectra of iWAT-adipocytes (ca. 100 adipocytes per mice per postnatal week, N=5 per measurement point) from three different spectral ranges. From the CH_2_ vibration region (2932-2778 cm^−1^) and the CO vibration region (1128-1080 cm^−1^), dynamic spectral changes were observed in iWAT-adipocytes at postnatal weeks 2-4 (AUC of the mean spectra decreases) and weeks 4-6 (AUC of the mean spectra increases). From the amide I and II bands (1700-1500 cm^−1^), an opposite direction of dynamic spectral change was observed compared to the other two regions. Additionally, we also observed how spectral features change between postnatal weeks 2 and 4, reflecting a transition from eWAT-like to iBAT-like spectra. In contrast, changes between postnatal weeks 4 and 6 reflect a transition in the opposite direction, i.e., from iBAT-like to eWAT-like spectra. To better understand the spectral dynamics for iWAT during postnatal weeks 2-6 in comparison to eWAT and iBAT, in **Fig.2i**, we show the kernel density estimation plot of PC1 where a clear spectral transition following tissue browning and tissue whitening is observed.

In order to quantify the iWAT transitions during postnatal weeks 2-6, we developed a metric based on logistic regression trained using depth-selective OA spectra of adipocytes from eWAT and iBAT (see **Methods**). In this way, we generated WAT- and BAT-scores for each adipocyte to indicate the similarity of spectra features to those from eWAT and iBAT in terms of a probability. Based on the WAT- and BAT-scores, we analyzed iWAT-adipocytes during postnatal weeks 2-6 in terms of their similarity to eWAT and iBAT, see **Fig.2j**. Thereby, we observed that the number of iWAT-adipocytes which is more similar to eWAT decreases from postnatal weeks 2 to 4 and reversely increases from postnatal weeks 4 to 6, while adipocytes spectrally more similar to iBAT increase from postnatal weeks 2 to 4 and decreases from postnatal weeks 4 to 6. **Fig.2j** represents trajectory of tissue browning (postnatal weeks 2-4) and the re-whitening (postnatal weeks 4-6) of iWAT-adipocytes based on WAT and BAT scores. Hence, adipose tissue remodeling can be analyzed based on the spectral features of adipocytes contained in iWAT by comparison with eWAT- and iBAT-adipocytes. Thus, depth-selective OA spectral analysis with spectral scoring can be used to detect and quantify the browning (postnatal weeks 2-4) and the re-whitening (postnatal weeks 4-6) phase of iWAT during postnatal development.

### Digital scoring of tissue based on MiROM and machine learning approach

Since the WAT and BAT scores were derived based on the spectra acquired from selected structures of interest, the analysis potentially suffers from a selection bias of manually chosen tissue locations, which could be corrected by performing hyperspectral analysis on the tissues. However, due to the acquisition time required for hyperspectral imaging using MiROM (e.g., 1×1 mm^2^ at step size of 5 μm for 502 wavenumbers will require ∼67 hours), the analysis was restricted to a localized spectral analysis of the chosen structures of interest. Nevertheless, to avoid selection bias and obtain spatial information (cell-to-cell variations) on adipose tissue remodeling, we extended our tissue scoring method and developed a quantitative spatial analysis tool (Q-SAT) for adipose tissue. Q-SAT allows to digitally score adipocyte features based on hyperspectral imaging with a reduced number of wavelengths compared to full spectral analysis to lower the data acquisition time (e.g., 1×1 mm^2^ (step size of 5 μm) requires only 80 minutes over 10 wavenumbers). In that manner, Q-SAT enabled the unbiased assessment of spatial distribution of adipose tissue remodeling at whole tissue level.

**Fig.3a** depicts the flowchart of Q-SAT. The underlying method of Q-SAT consists of a logistic regression model trained on the depth-selective OA signal intensities of 1) adipocytes from eWAT and iBAT, and 2) further tissue structures located in the iWAT, including ECM, lymph nodes, connective tissue, water, etc. The model segments and scores adipocytes with respect to spectral features related to WAT or BAT based on 10 wavenumbers (2856, 1632, 1550, 1454, 1376, 1238, 1134, 1086, 1046, 990 cm^−1^), selected after analyzing spectra obtained on each tissue structure, see **Methods** for details. The trained model was then applied to segment and digitally score adipocyte features in hyperspectral iWAT images. Similar to the previously discussed tissue scoring, the digital scoring is related to adipocyte features that are either corresponding to WAT or BAT. **Fig.3b** shows an overlay of the OA absorption map of iWAT based on two vibrational contrasts (yellow: 2856 cm^−1^, cyan: 1550 cm^−1^) at the postnatal week 6 from a large FOV (5×10 mm^2^). Below the OA absorption map, a digital scoring map of the same FOV is visualized (**Fig.3c**). To monitor tissue remodeling in iWAT during postnatal development, we applied Q-SAT to images from all postnatal stages of iWAT in a small FOV (1×1mm^2^), as shown in **Fig.3d,** and plotted the digital contrast in more intuitive way in **Fig.3e, f**. to quantitatively compare WAT and BAT scores produced by Q-SAT during postnatal development (3 mice per postnatal week). Thereby, as expected, we observed that the relative overall WAT score of iWAT decreased during the browning phase of postnatal weeks 2-4, while the relative overall BAT score increased. The inverted trend was observed during the whitening phase of the postnatal weeks 4-6. With Q-SAT we unveiled the spatially heterogeneous appearance of adipose tissue remodeling while simultaneously avoided the selection bias of localized spectral analysis associated with manual structure selection.

**Figure 3.**
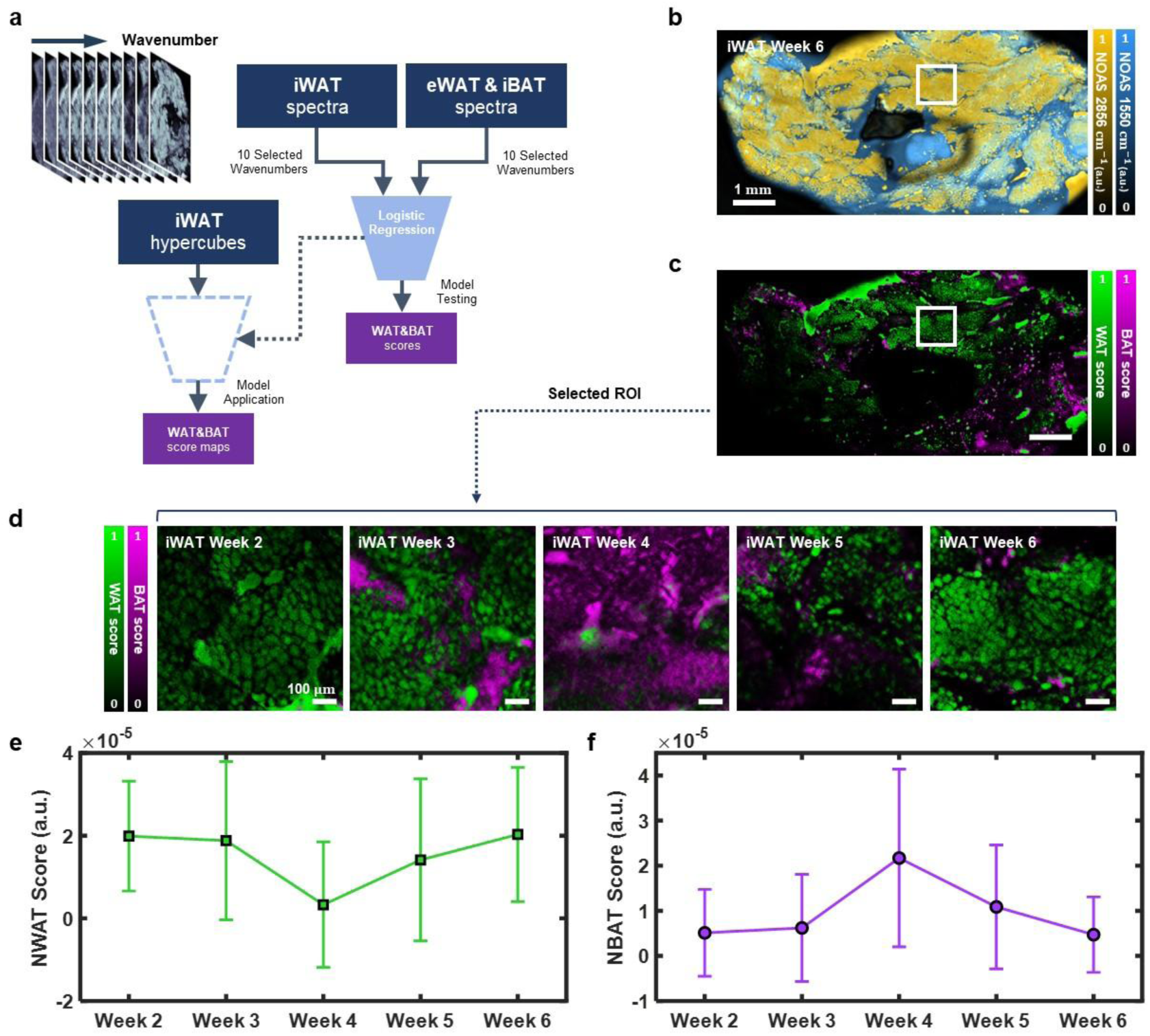
Quantitative spatial analysis of postnatal tissue remodeling in inguinal white adipose tissue. **(a)** Flowchart of the quantitative spatial analysis tool (**Q-SAT**) used for pixel-wise digital scoring the level of WAT or BAT similarity of iWAT. **(b)** Merged depth selective OA micrograph at 2856 *cm*^−1^ (yellow) and 1550 *cm*^−1^ (cyan) for a large FOV (5×10 *mm*^2^) collected during iWAT postnatal week 6. **(c)** WAT and BAT score mapping results for **b**. **(d)** Digital scoring map for a 1×1 *mm*^2^ FOV area during postnatal development (weeks 2-6) in iWAT. **(e, f)** Mean normalized WAT and BAT (**NWAT**, **NBAT**) scores for the 1×1 *mm*^2^ FOVs images shown in d. The error bars indicate standard deviation. Results in d, e obtained from three independent measurements (N=3) per postnatal week.

## Discussion

In this study, we demonstrated that MiROM can quantitatively monitor the intracellular changes of biomolecules in fresh unprocessed fat tissues while preserving the native tissue architecture in a depth-selective and label-free manner. In particular, intracellular multi-spectral imaging and spectral analysis using MiROM enabled comparison and differentiation of the morphological and molecular hallmarks of eWAT and iBAT, as well as the monitoring of dynamic WAT remodeling. We thus developed a new methodology that can aid in the identification and characterization of adipose tissue plasticity and heterogeneity, providing a powerful tool for studying tissue physiology and pathology in situ.

With MiROM, we observed distinctive morphological and molecular differences between adipocytes in eWAT and iBAT. For instance, as expected from H&E-based histology, we observed that eWAT-adipocytes are ca. 3.3 times larger than iBAT-adipocytes and that the lipid contrast (at 2856 cm^−1^) from eWAT-adipocytes is ca. 1.7 times higher than the lipid contrast originated from iBAT-adipocytes.^38^ Furthermore, as expected from western blot measurements, we observed that the protein contrast (at 1550 cm^−1^) in iBAT-adipocytes is ca. 3.4 times higher than in eWAT-adipocytes. This confirms that, compared to iBAT, eWAT is a lipid-rich and protein-poor tissue, mostly due to its larger size of lipid droplets and a lower number of mitochondria, as shown in previous studies on mitochondrial abundance and lipid droplet morphology.^38^ Additionally, by analyzing the intrinsic molecular and morphological hallmarks of eWAT and iBAT obtained with MiROM, we were able to monitor iWAT browning during postnatal weeks 2 to 4 in mice and its subsequent re-whitening during postnatal weeks 4 to 6. This observation was confirmed by western blot, H&E staining, and whole-mount immunofluorescence staining (**Supplementary Fig.4**). Moreover, beyond qualitative imaging, we assessed the spatial distribution of adipocyte types in different postnatal weeks during adipose tissue remodeling process and observed substantial heterogeneities with respect to different postnatal weeks based on pixel-wise tissue scoring with Q-SAT. Since Q-SAT uses only a low number of spectral points (10 wavenumbers), it significantly increased the hyperspectral image acquisition speed (50-fold acceleration). Collectively, MiROM offers technology for intracellular analysis of adipose tissue in a label-free and non-destructive manner by enabling the assessment of the native tissue architecture and intrinsic biomolecule while avoiding time-consuming tissue preparation, including fixation and staining.

Here, focused on characterizing overall lipid and protein content in adipocytes during adipose tissue remodeling. However, since lipid composition and the number of mitochondria play a critical role in the metabolic function of adipocytes, for a better understanding of adipose tissue remodeling, combining MiROM readouts with mass spectroscopy-aided in-depth lipidomics and proteomics could reveal the sample’s single-biomolecular composition. Additionally, we anticipate that the combination of hyperspectral imaging using MiROM aided with Q-SAT and mass spectrometry cross-validation will provide a quantitative approach for assessing biomolecules of adipocytes while providing native morphological information. Furthermore, while Q-SAT significantly reduced the required number of spectral points for the assessment of the spatial distribution of adipocyte types, the hyperspectral imaging acquisition time is still limited by the mechanical properties of MiROM, i.e., the speed and acceleration of the motorized stages used for point-by-point raster scanning. To address the limitations associated with long imaging times, new strategies to achieve fast image acquisition speed are currently being implemented in our lab. These involve galvo- or MEMS (micro-electro-mechanical systems)-mirrors, simultaneous acquisition of OA transients with multiple wavenumbers, increasing laser repetition rate up to 2MHz, and sparse data acquisition combined with Bayesian image reconstruction.^39^

In summary, we characterized molecular hallmarks of eWAT and iBAT and monitored intracellular molecular and morphological changes of adipocytes during postnatal transient adipose tissue remodeling using MiROM. As a result, we identified novel biochemically distinguishable characteristics between WAT and BAT, developed a new methodology that can aid in the identifying and monitoring of the adipose tissue browning and whitening process and gained insights into the metabolic plasticity of adipose tissue at the intracellular level. With our AI-based quantitative spatial tissue analysis tool (Q-SAT), we further obtained spatial assessments of beige adipocyte development at the tissue level. In the future, the combination of MiROM and Q-SAT will be expanded to examine further applications related to adipose tissue (e.g., margin assessment of fatty tumors), leveraging the high precision of fast, label-free histological assessments for its integration into clinical workflows.

## Methods

### MiROM configuration and measurement process of adipose tissue

MiROM in transmission mode has been shown in **Fig.1a**. A pulsed QCL (MIRcat, Daylight Solutions) was used as a mid-IR excitation source for OA signal generation. Since the QCL spectral range covered 3.4-11 μm (2932-909 cm^−1^) and the full width at half maximum (FWHM) of spectral linewidth was <1 cm^−1^, biomolecular specificity was able to be achieved during the measurement. The focused mid-IR laser beam (Repetition rate: 100 kHz, pulse duration: 20 ns), focused by 0.5 NA reflective objective (36×, Newport Corporation), excited the sample located on top of the mid-IR transparent ZnS window (Crystal) of the custom-designed metallic petri dish. The tissue sample was placed on the ZnS window after removing the superficial buffer solution from the surface of the tissue to decrease the mid-IR attenuation, which is driven by the strong mid-IR absorption of water. An acoustically transparent plastic membrane covers the tissue with ultrasound gel to isolate the tissue from the coupling media (Deionized water) while it is feasible to transfer the generated acoustic wave from the focal spot of the mid-IR beam. The generated OA signal has been acquired by a focused ultrasound detector (Imasonic and Sonaxis), which has 20 or 25 MHz central frequency, immersed in coupling media. The raw OA signals were amplified with a 63 dB low-noise amplifier (MITEQ) and filtered with a 50 MHz low pass filter (Mini-circuits). Filtered OA signals recorded by 200 MS/s on a data acquisition (DAQ) card (Gage Applied). To avoid the interference of CO_2_ and water vapor, the optical beam path has been covered by the purged inert gas chamber with dry N_2_ gas.

The lateral and axial resolution of MiROM has been experimentally assessed by the measuring point-spread-function (PSF) of a polystyrene sphere that size of 1 μm at 2850 cm^−1^(Lateral resolution: 5.3 μm, axial resolution: 42.2 μm). The detailed optical performance of MiROM is reported elsewhere^25^.

### Depth-selective analysis and lateral segmentation of adipocytes

To compare the interior characteristics of adipocytes within the same depth range, we utilized a time dependency of the OA signal. Since the acoustic wave allows us to estimate the point of signal generation from signal arrival time, we could deduct axial information of adipocytes. To analyze the OA signal at the specific depth range, we acquired the absolute (Abs.) of the OA analytic signal from the OA transient (A-line). All OA analytic signals were obtained from the OA transients, which are interpolated by cubic interpolation. We defined the minimum size of the time window (depth range) as 7.5 μm (1 sampling time point; sampling rate: 200 MS/s, estimated acoustic wave speed at soft tissue: 1500 m/s). For the intracellular comparative analysis of each tissue, we acquired each depth-selective intensity of the mid-IR absorption map (micrograph) and spectra based on the area under the curve of the Abs. of OA analytic signal from the defined time window. Since the primary content of adipocytes is lipid droplet, we defined an intracellular time window based on maximum amplitude depth in 2856 cm^−1^.

Additionally, to localize adipocytes from extracellular components, we also laterally segmented mid-IR absorption map through intensity thresholding. We obtained segmentation masks from each micrographs with the pixels with intensities higher than the first quantile (25% of data) of iBAT’s OA contrast at 2856 cm^−1^.

### MiROM image acquisition and processing

Since the system was designed based on the co-alignment of the mid-IR excitation beam and focused ultrasound detector, to obtain the mid-IR absorption map, the OA transients have been acquired by point-by-point raster scanning pattern of x-y motorized stage (Physik Instrumente). The optical and acoustical focal planes have been adjusted by the z-axis mechanical stages, and the ultrasound detector and reflective objective are individually mounted. Each pixel intensity of the micrographs has been obtained by the chosen wavenumbers from 2932-2770 cm^−1^ and 1738-909 cm^−1^ at the corresponding position assigned by the x-y motorized stage. We acquired intensity in two ways from the OA transient: peak-to-peak intensity of OA transient (MAP) or depth-selective intensity, which is mentioned in the previous section. At each pixel position, 50 OA transients have been acquired (since the pulse repetition rate is 100 kHz, the pixel acquisition time is 500 μs.). The averaged intensity of 50 OA transients has been formed as an intensity of each pixel. Micrographs are obtained at a FOV of 500×500 μm^2^, 1×1 mm^2^ with 2 or 5 μm step sizes (500×500 μm^2^ (step size – 2 μm): 13 minutes, 1×1 mm^2^ (step size – 2 μm): 40 minutes, 1×1 mm^2^ (step size – 5 μm): 8 minutes per single wavenumber).

For the contrast enhancement of the micrograph, to analyze the structural details, two image post-processing steps have been utilized on all MiROM micrographs in the figures displayed in the manuscript: normalized square intensity of the OA signal (OAS), 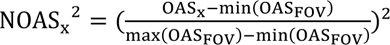, and contrast limited adaptive histogram equalization (CLAHE). Processed micrographs acquired from 2856 cm^−1^have been utilized to analyze the size of adipocytes from each adipose tissue, respectively. The adipocyte boundaries were defined based on the contrast difference between adipocytes and extracellular structure of adipocytes at 2856 cm^−1^, and the size of adipocytes was calculated based on the Cell Profiler 4.0 software.

### MiROM spectra acquisition and processing

The mid-IR absorption spectra have been obtained while tuning pulse tunable QCL with step size 2 cm^−1^from 2932-2770 cm^−1^and 1738-909 cm^−1^at corresponding localized locations from the sample. To enhance the accuracy of the spectral analysis of adipocytes, 10000 OA transients per wavenumber at each adipocyte have been obtained to increase the signal-to-noise ratio (SNR). The spectral acquisition time of each adipocyte is ∼8 minutes. The raw OA intensity profile across wavenumbers obtained at MiROM spectral analysis, displayed in the manuscript, was depth-selective intensity.

Since the OA signal has been affected by the daily response of the system, such as laser fluctuation, alignment of optics and acoustic detector, to compensate for the slight variations on the laser emission profile of the QCL, we obtained OA spectral intensity across wavenumbers by correcting the sample’s raw OA intensity profile (Intensity: depth-selective intensity) dividing by OA intensity profile (Intensity: peak-to-peak value of OA transient) of the reference sample (carbon adhesive tape). Since the carbon adhesive tape shows broadband absorption at the emission range of the QCL, it allows the measurement of the laser emission profile of the QCL.

### Machine-learning-based digital staining and tissue scoring

Combining and generating a metric to describe the spectral hallmarks of BAT and WAT enables quantitative tracking of the tissue remodeling presented in this work. To exhibit the temporal development of browning and whitening of iWAT, we, therefore, implemented a machine-learning approach to quantify tissue properties based on spectral information. The workflow used for modeling was to train a logistic regression model based on the reference spectra extracted from eWAT and iBAT while labeling the tissue types (WAT&BAT) of the corresponding spectra. Applying the trained model to the spectra collected from adipocytes of individual postnatal stages and extracting the predicted probabilities for WAT&BAT allows us to statistically assess the predicted probabilities used to score spectral properties over the postnatal stages. In a subsequent approach, we selected 10 wavenumbers as model features determined using PCA containing information about the spectra’s corresponding tissue types and trained another logistic regression model based on the selected wavenumbers to ultimately apply the model to hyperspectral tissue images for digital staining. Therefore, we additionally included spectra from iWAT corresponding to additionally labeled structures apparent in the images, such as ECM, lymph nodes, water, and void area, to enable discrimination of the model between spectral hallmarks related to adipocytes and other structures. Applying the trained model to the hyperspectral images acquired based on the 10 excitation wavenumbers and extracting the pixel-wise prediction probabilities enables digital staining of spectral hallmarks related to BAT and WAT in a spatial context. Similar to the statistical evaluation of the spectral scoring, the scores from the images taken from iWAT were statistically assessed over all pixels to quantify the overall tissue score and thus track the tissue remodeling process over time. The overall metric for the normalized tissue scores *S*_WAT_ and *S*_BAT_ extracted for each hyperspectral image in every postnatal week was computed based on the pixel-wise model outputs WAT(𝑥, 𝑦) and WAT(𝑥, 𝑦) integrated over the entire image, i.e.,

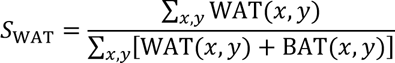

and

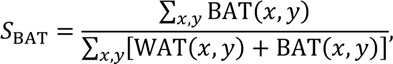

to put the WAT&BAT scores concerning the overall content related to structures with spectral hallmarks of adipocytes. The normalized tissue scores *S*_WAT_ and *S*_BAT_ enable digital staining of spectral properties related to WAT&BAT and thus constitute a statistical metric to track the tissue remodeling process.

### Feature extraction and data preprocessing

Tissue scoring requires features to be extracted from OA signals and preprocessed for training and application of the machine learning model. The raw signals acquired using MiROM are OA transients carrying structural information about the absorption coefficients in the sample. The digital staining model uses depth-selective intensities as features derived from the A-lines according to the ‘Depth-selective analysis and lateral segmentation of adipocytes’ section. The extracted intensity values are then divided by the corresponding carbon spectrum intensities to compensate for the laser emission profile. For training the spectral scoring, the spectra were normalized to the uniform spectral norm by dividing each spectrum 𝐬 = [𝐼(𝜆_0_) ⋯ 𝐼(𝜆_𝑁_)]^T^ composed by OA spectral intensity values for each wavenumber 𝜆 by the spectral L2-norm, i.e.

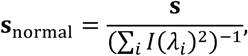

to remove intensity drifts in the system and only account for the spectral shape to determine the scoring. While 𝑁 = 502 for the spectra taken at adipocytes, the normalization was applied analogously to the 10-wavenumber hypercubes for each datapoint (pixels in the hyperspectral images) individually, where 𝑁 = 10. In order to ensure convergence and high accuracy of the algorithm to minimize the objective function given by the categorical cross-entropy of the training dataset, the features were scaled using standardization, i.e.

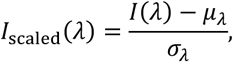

where 𝜇_𝜆_ are the mean and 𝜎_𝜆_ the standard deviation of all intensities for the wavenumber 𝜆 contained in the training dataset. Scaling unseen data, which are not part of the training dataset, was done analogously using 𝜇_𝜆_ and 𝜎_𝜆_ derived from the training data.

### Adipoclear

Whole Tissue Clearing was performed following the Adipo-Clear method published by Chi et al. in 2018.^40^ In short, whole iWAT pads were fixed with 4% paraformaldehyde in PBS for 24 hours at room temperature. Fat pads were dehydrated using a series of methanol gradients in glycine buffer. Fat pads were delipidated in 100% dichloromethane, washed with 100% methanol, and subsequently bleached in a 5% H_2_O_2_ solution in methanol. After rehydration in a reverse methanol/glycine buffer gradient, fat pads were stained against UCP1 (Abcam, ab23841) at a 1:200 dilution containing Heparin and Triton X-100 for one week. Fat pads were then washed and incubated for another week with a multi-rAb CoraLite Plus 647-Goat Anti-Rabbit 647 secondary antibody (Proteintech, RGAR005) at a 1:500 dilution. Lint was gently removed from the fat pads under a dissection microscope. Fat pads were then embedded in a 1% Agarose in PBS solution to stabilize its morphology. Embedded fat pads were again dehydrated in a methanol gradient and delipidized in 100% DCM. Samples were incubated overnight in ethyl cinnamate for refractive index matching and analyzed by fluorescent microscopy (Zeiss, LSM 700).

### Histology

iWAT was isolated and immediately fixed in 4% paraformaldehyde for 24 hours at room temperature. Tissues were then dehydrated using an ethanol gradient and embedded in paraffin. Tissues were sectioned into 5 μm slices and stained with hematoxylin and eosin. Slices were analyzed using an EVOS XL Core Imaging System.

### Tissue Preparation for MiROM imaging

Whole adipose tissue pads were carefully isolated from FVB/NJ mice. eWAT and iBAT pads were isolated from 6-week-old mice, while iWAT pads were isolated from mice aged between 2-6 weeks. Tissues were lightly fixed for 4 hours in IC fixation buffer (eBioscience, 00-8222-49) at room temperature. After fixation, tissues were stored in PBS at 4°C until the day of measurement.

### Western Blotting

Whole iWAT pads were isolated and snap-frozen in liquid nitrogen. For protein isolation, 100-200 μl RIPA buffer was added to each whole fat pad. Fat pads were homogenized for 20 seconds using an UltraTurax T10 (IKA). The resulting homogenate was centrifuged at 4°C for 10 minutes at 14.000 g. The supernatant was collected, and protein concentration was measured using the Pierce BCA Protein Assay (Thermo Scientific, 23225). 30ug of Protein was loaded and then separated on a 12.5% PAGE Gel. Proteins were blotted onto a nitrocellulose membrane for 1h at 100 mA using a SemiDry blotting approach (Analytik Jena, Biometra Fastblot)). The membrane was blocked with 5% BSA for 90 minutes at room temperature and then incubated in a 1:2000 rabbit anti-UCP1 (Abcam, ab23841) and 1:5000 mouse anti-Beta-Actin (Proteintech, 66009-1-Ig) primary antibody solution overnight at 4°C. After multiple washes, secondary antibodies against rabbit and mouse (LICOR, 926-32211 and 926-68070) were added at 1:20000 dilutions for 90 minutes at room temperature. Protein signals were detected using the 700 nm and 800 nm channels of a LICOR Odyssey FC Imager.

## Acknowledgements

This work was supported by DFG Research Unit iMAGO-FOR5298 (455422993) to Y.L., M.A.P., and Ma.K., DFG-TRR 333/1 – 450149205, Emmy Noether program (441904031) and ERC starting grant (101078516) to Y.L., and from the European Research Council (ERC) under the European Union’s Horizon 2020 research and innovation programme under grant agreement No 694968 (PREMSOT) awarded to V.N. We thank Dr. Serene Lee and Dr. Elisa Bonnin for their attentive reading and improvements of the manuscript.

## Contributions

M.K., A.W., Y.L, and M.A.P conceived and designed the study. M.K., A.W., and A.P. performed the experiments. M.K. and C.B. performed spectral data analysis. M.K., A.W., and C.B. prepared the figures. C.B. developed digital scoring method. A.W. performed validation measurement of western blot, H&E staining, and whole-mount immunofluorescence staining. V.N. provided support on optoacoustic detection. Ma.K. provided critical resources related to mouse maintenance and tissue preparation. M.K., A.W., C.B, Y.L., and M.A.P wrote the manuscript. Y.L. supervised tissue acquisition and remodeling experiments. M.A.P supervised the acquisition of OA images and spectra on mice tissues. Y.L. and M.A.P. supervised the whole study. All authors read and edited the manuscript.

## Competing interests

V.N. and M.A.P. are founders and equity owners of sThesis. V.N. is a founder and equity owner of Maurus OY, iThera Medical GmbH, Spear UG, and I3 Inc. The other authors declare no competing interests. A patent application (WO 2019 149 744 A1) licensed to sThesis GmbH, relevant to the technology discussed in this paper, has been filed.

## Supplementary Figures

**Supplementary Figure 1.**
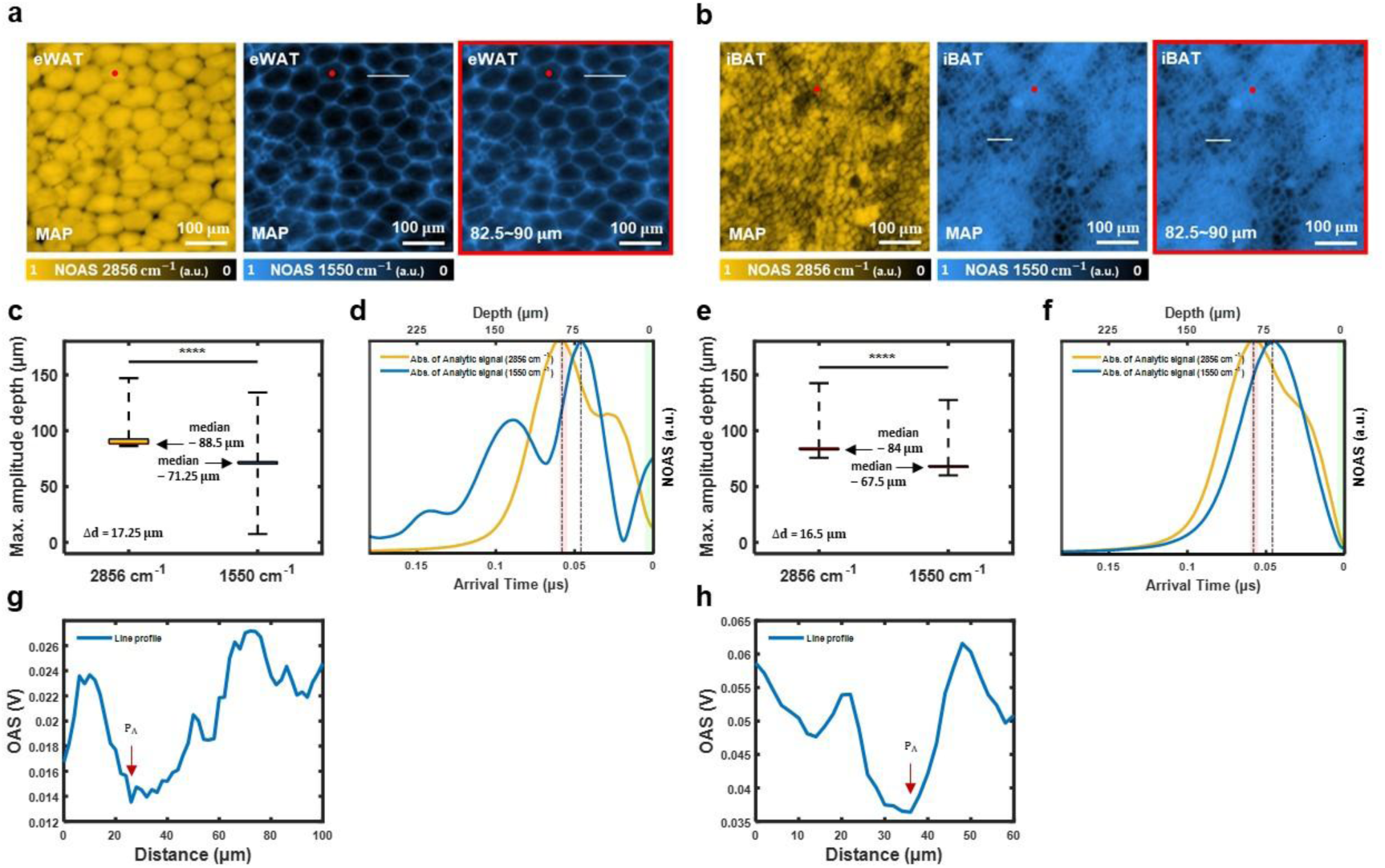
Comparison between maximum amplitude projection (MAP) analysis and the depth-selective analysis on eWAT by MiROM. **(a, b)** MAP and depth-selective OA micrograph of eWAT and iBAT at 2856 and 1550 cm^−1^ channel. Depth-selective OA micrograph pixel intensity represents the area under the curve of the Absolute (**Abs.**) of the OA analytic signal within the selected depths. **(c, e)** Box plot showing the MAD at 2856 cm^−1^ and 1550 cm^−1^. Asterisks (****) denote the level of statistical significance (P<0.0001) analyzed by a two-sample t-test (**c**: eWAT, **e**: iBAT). **(d, f)** Abs. of OA analytic signal from 2856 and 1550 cm^−1^ (**d**: eWAT, **f**: iBAT). The red box indicates the time window (82.5∼90 μm), which contains the depth of interest (MAD at 2856 cm^−1^; width: 7.5 μm). **(g, h)** Contrast line profiles, white line from at MAP OA micrographs in 1550 cm^−1^ (**g**: eWAT MAP OA micrograph, **h**: iBAT MAP OA micrograph).

**Supplementary Figure 2.**
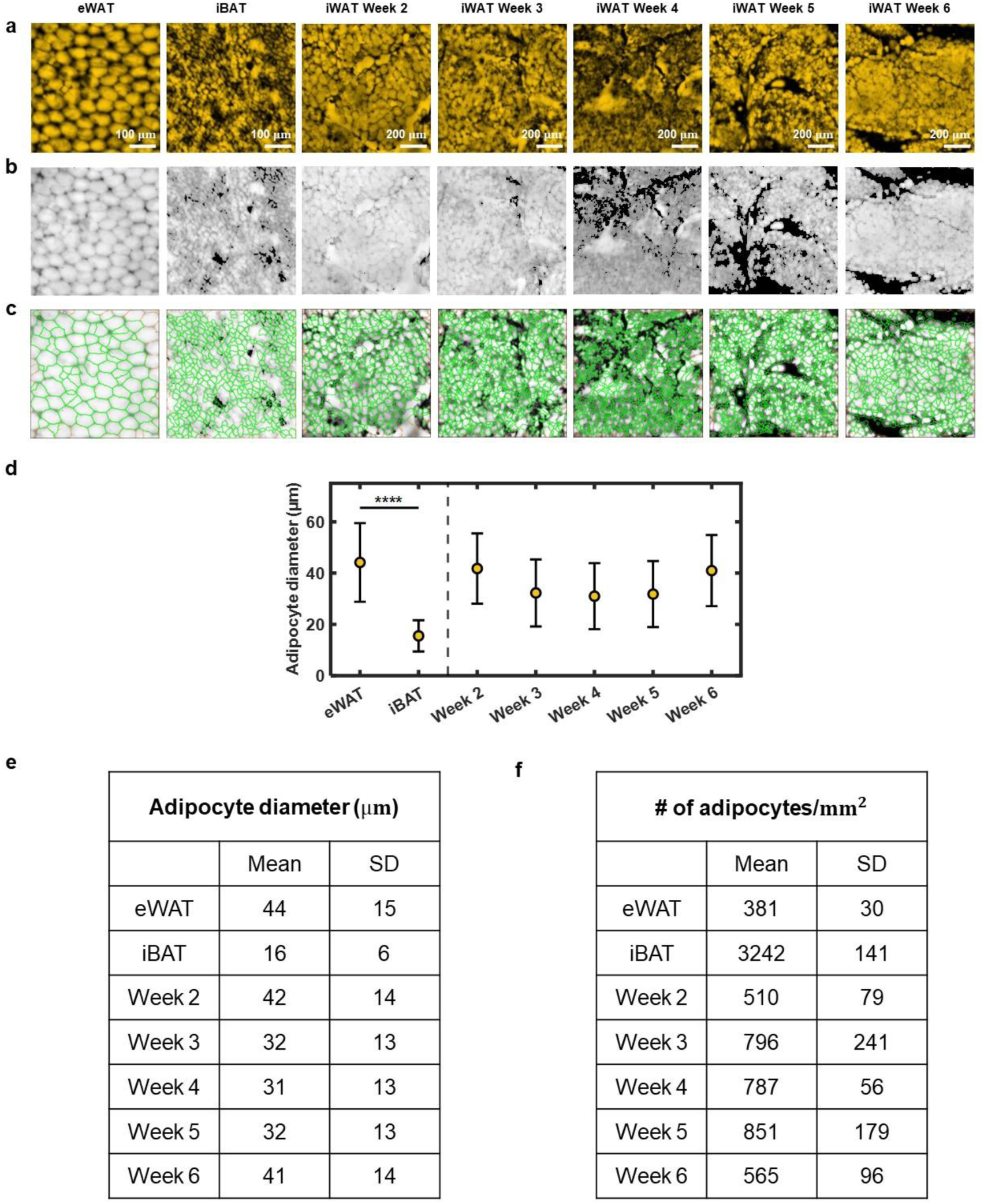
Intensity threshold-based masking results and adipocyte diameter and adipocyte density (adipocyte count per unit area) calculation at depth-selective OA micrograph. **(a)** Depth-selective OA micrograph of eWAT, iBAT, and iWAT of postnatal weeks 2-6 at 2856*cm*^−1^ channel. **(b)** Intensity threshold-based masking results of depth-selective OA micrographs of eWAT, iBAT, and iWAT of postnatal weeks 2-6 at 2856*cm*^−1^ channel. Lateral segmentation based on intensity thresholding highlights the location of adipocytes. **(c)** Adipocyte segmentation produced by Cell Profiler 4.0 at selected eWAT and iBAT regions of interest (**ROI**). **(d)** Mean adipocyte size from ROI of eWAT, iBAT, and iWAT of postnatal weeks 2-6 (eWAT and iBAT: N=7 for each tissue type; iWAT at postnatal weeks 2-6: N=5 for each tissue type) with error bars indicating standard deviation (**SD**). Asterisks **** denote a statistically significant difference (P<0.0001) between eWAT and iBAT, and it has been analyzed by a two-sample t-test. **(e, f)** Mean and standard deviation of adipocyte diameter and adipocyte count per unit area (*mm*^2^) of eWAT, iBAT, and iWAT of postnatal weeks 2-6.

**Supplementary Figure 3.**
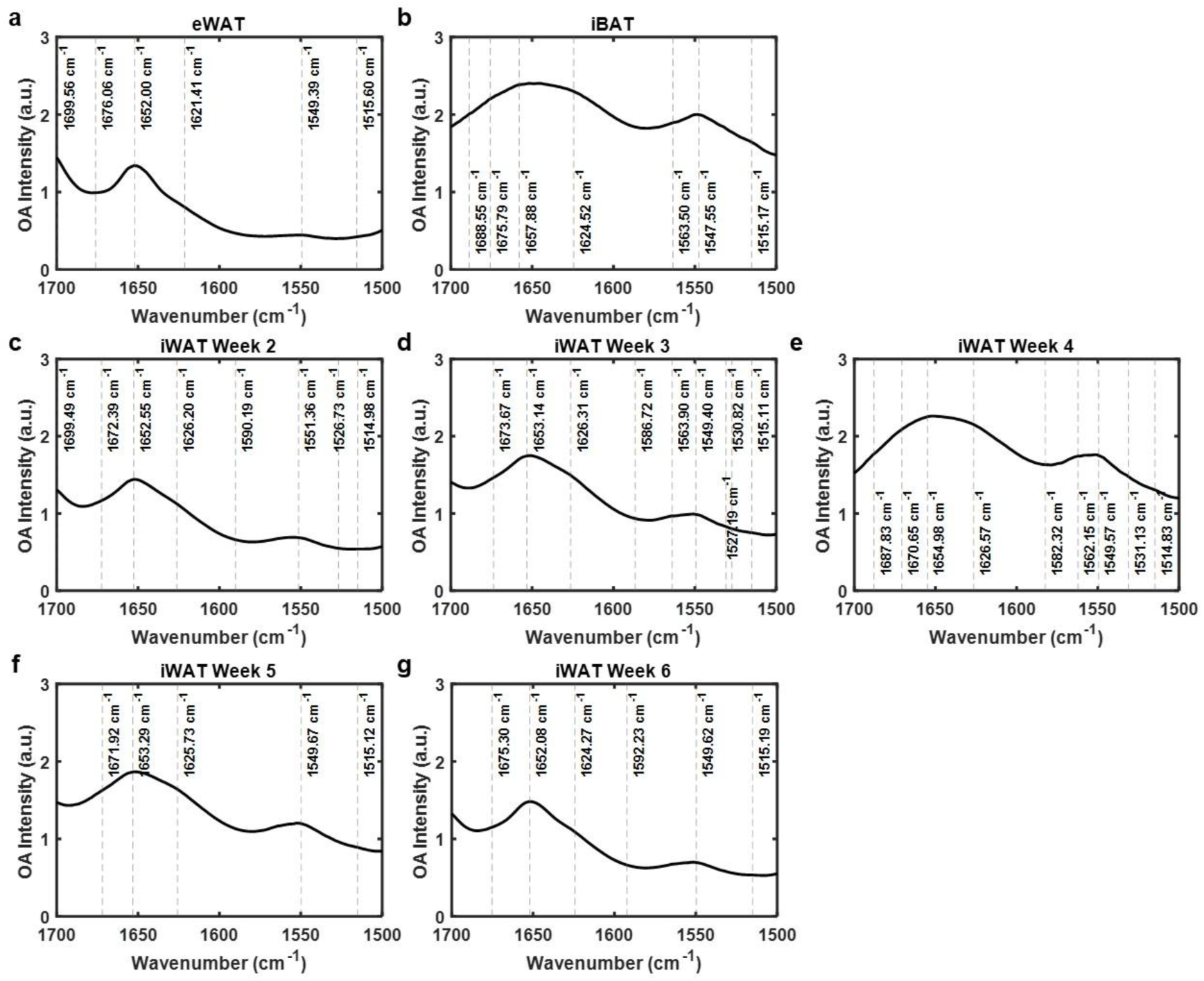
OA spectra at 1700-1500 𝐜𝐦^−𝟏^. Mean OA spectra (solid black line) of eWAT, iBAT, and iWAT of postnatal week 2-6 are shown. With a dashed line, local minima points from the second derivative spectra, which indicate vibrational frequency associated with biomolecules, are shown. The amide I and II bands were analyzed to retrieve spectroscopic features of intrinsic biomolecules. **(a)** eWAT, **(b)** iBAT, and **(c-g)** iWAT in postnatal stages (postnatal weeks 2-6). eWAT and iBAT spectral features show a distinctive difference. iWAT spectral feature transition between postnatal week 2 and 4 reflects the transition from eWAT-like to iBAT-like spectra, and the transition between postnatal week 4 and 6 shows the opposite direction.

**Supplementary Figure 4.**
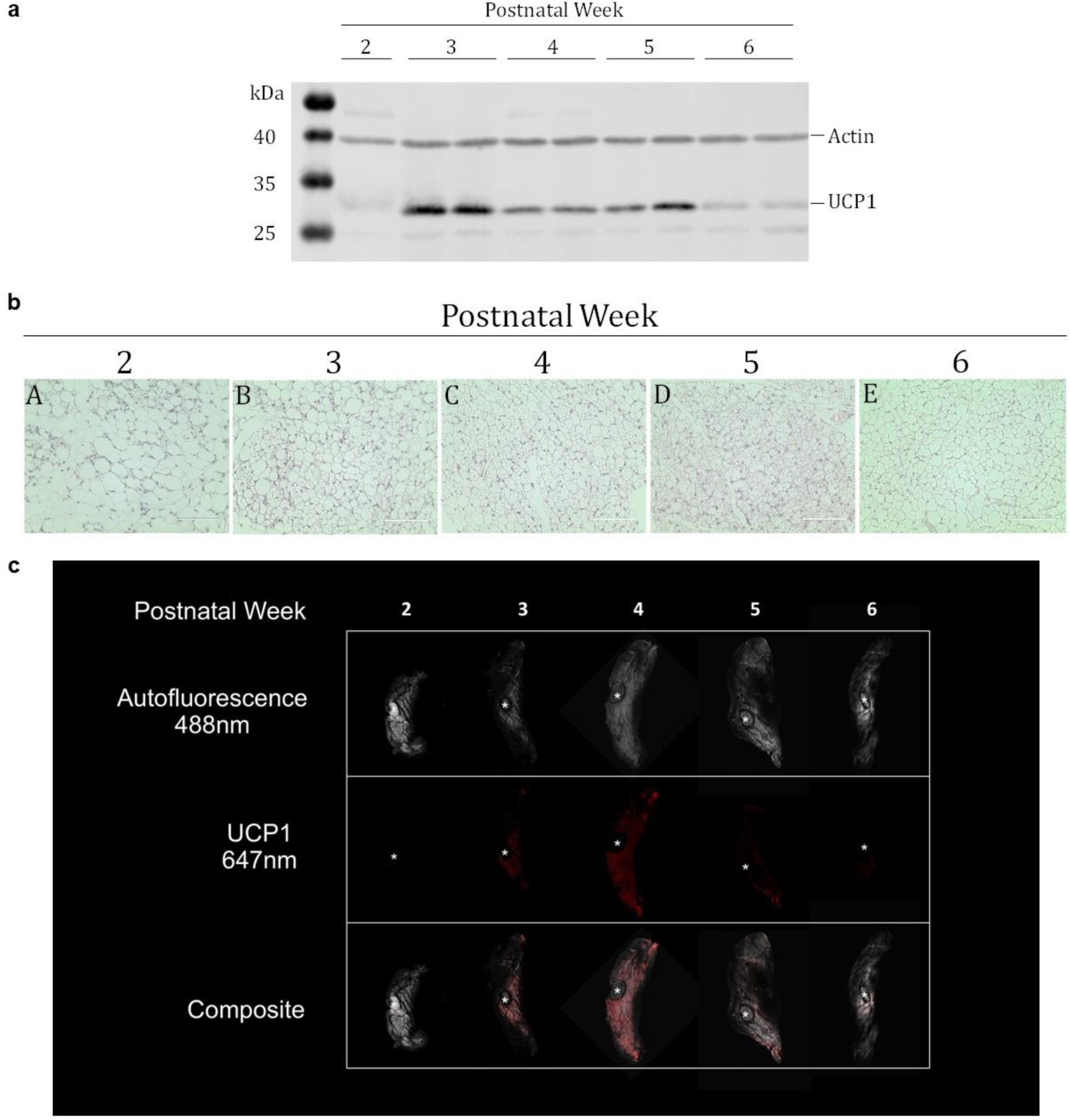
Biological validation of uncoupling protein 1 (UCP1) levels and morphological changes of adipocytes are transiently elevated in iWAT during postnatal development. **(a)** Protein was isolated from iWAT during postnatal weeks 2-6. Immunoblots were stained against UCP1 (Abcam, ab23841) and beta-Actin (Proteintech, 66009-1-Ig). **(b)** Histological sections of iWAT from postnatal weeks 2-6 stained with hematoxylin and eosin (**H&E**) and depicted at 40x magnification. All sections were captured near the LN present in iWAT to ensure comparability between histological sections and MiROM measurements. **(c)** The panel depicts iWAT from postnatal weeks 2-6. Each image is a stitched optical section captured with a 5x objective. Each column depicts one iWAT sample with its autofluorescence in grey, an immunolabel against UCP1 in red, and their composite. The LN of each tissue is marked by *.

## Notes

### Summary of Updates

Dear bioRxiv editorial office manager, We just realized we mis-indicated the wrong funding number at the funding information. Therefore, we changed the funding information at the manuscript and the system. Rest of the information is the same as before. Thank you for your kind concern. Best regards, Myeongseop Kim

